# Neural networks learn forward dynamics when freed from numerical integration

**DOI:** 10.64898/2026.05.27.728310

**Authors:** Serhii Bahdasariants, Sergiy Yakovenko

## Abstract

Seamless interaction between humans and machines requires interfaces that remain robust to the variability inherent in biological signals and physical environments. Advanced human-machine interfaces (HMIs) increasingly rely on machine learning to predict or control limb dynamics. These systems must learn input-to-output mappings between control variables and limb state, such as the mapping from muscle forces or joint torques acting about segmented arm joints to limb posture over time. Such statistical input-to-output transformations can result in numerical instability of predicted musculoskeletal kinematics and dynamics. Achieving the robustness of biological motor control requires solving both forward and inverse dynamics problems; however, these problems are computationally asymmetric because they entail opposing operations, integration and differentiation. Since we have previously shown that neural networks solve the inverse dynamics problem when trained to map kinematic to dynamic signals during reaching, we hypothesized that representing separately the approximation of equations of motion (EOM) and their temporal numerical integration may capture the relevant computational structure of the forward dynamics problem. We tested this hypothesis by comparing a conventional direct-mapping neural network with a two-stage model, the artificial physics engine (APE). When predicting the state of a two-segment system under external perturbations not encountered during training, the direct-mapping, monolithic model produced large prediction errors inconsistent with the expected interaction torque, whereas the APE maintained low error and remained stable under novel initial conditions and perturbations. Mapping system dynamics in terms of the EOM imposes a causal, physics-based structure on forward models for HMI design.

## INTRODUCTION

The difficulty of modeling complex dynamics is not determined solely by the underlying physics, but also by the representation in which those dynamics are expressed. Those representations can differ markedly in compactness, stability, and computational tractability. For example, in biomechanics, the equations of motion (EOM) can be expressed with Newton-Euler, Lagrange, and Kane’s methods, among others, to capture the input–output relationships as state-evolution problems that require temporal integration (Featherstone, 2008; Yamaguchi, 2005). In control theory, the difference between instantaneous algebraic mappings (*u* → *x*) and state-evolution equations 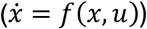 separates input-output models without explicit internal state from those with explicit evolution of internal system state (Kalman, 1963). Many machine-learning approaches to movement decoding are formulated in the former way, as direct mappings from measured signals to outputs, rather than as explicit models of state evolution (Glaser et al., 2020; Mathis et al., 2024). While the approximation of system dynamics improves with network depth and the sufficient representation of state behavior within the training dataset (Bengio, 2009), this representation may not be well-suited.

In our previous studies developing movement control algorithms for intuitive human-machine interfaces (HMIs), we have achieved real-time performance in arm and hand simulations with optimized musculoskeletal transformations (Sobinov et al., 2017) optimizing Hill-type muscle models with traditional polynomial and, later, artificial neural network (ANN) representations (Gritsenko et al., 2016; Smirnov et al., 2021; Sobinov et al., 2020). The segmental forward dynamics were computed using an optimized physics engine, MuJoCo (Todorov et al., 2012), and the decoding of electromyographic (EMG) signals into torques, based on the muscle intrinsic state and joint-angle-dependent moment arms, was used to map the EMG signals to the intended movement. This type of mapping can be captured with physics-informed neural networks (PINNs) that enforce governing physics equations directly in the loss function (Raissi et al., 2019). While PINN formulations can represent both forward and inverse dynamics by constraining differential equations (by minimizing explicitly stated loss terms derived from the governing physical equations), their optimization may become challenging for noisy, partially observed, nonstationary real-time biomechanics (Cuomo et al., 2022; Karniadakis et al., 2021). Encoding movement features with PINNs requires full-motion motifs, thereby limiting their applications to state prediction (Taneja et al., 2024). Alternatively, physics-informed recurrent dynamics, captured by recurrent neural networks (RNNs), are similar to causal integration and local state evolution in cortical (Lillicrap and Scott, 2013) and cerebellar dynamics (Miall et al., 1993; Miall and Wolpert, 1996), as well as the recursive estimation needed for the sensorimotor loop (Kording et al., 2004; Todorov and Jordan, 2002). However, while the corresponding motion equations for arm and hand inverse dynamics can be represented within even a shallow ANN (Manukian et al., 2023), without MuJoCo, the forward simulations of similar complexity have not yet been achieved using machine learning. Several prominent studies have demonstrated gesture-based control for up to 10 degrees of freedom (DOFs); however, the continuous robust decoding is typically limited to only 2-3 DOFs (Ajiboye et al., 2017; Collinger et al., 2013; Hochberg et al., 2012; Sussillo et al., 2016). Therefore, despite evidence that limb-dynamic computations can be embedded even in shallow neural networks, producing physics-constrained forward online inference remains challenging.

In this study, we test the hypothesis that simulation instability in forward dynamic transformation arises from a monolithic ANN implicitly conflating the approximation of the EOM with the computationally demanding process of numerical integration. We implement the artificial physics engine (APE), a decoupled physics-informed recurrent forward-dynamics architecture composed of two distinct modules: (1) an ANN trained *only* to approximate the terms in the EOM (a forward dynamical mapping), and (2) a dedicated, external numerical integrator that handles the state update. We used perturbations that were not included in the training as evidence of learning not only the input-output mapping but also embedding body mechanics, with interaction torques propagating along the segmented structure. Evidence of embedded physics in neural computations opens opportunities to formulate accurate and adaptable internal models of body structure for HMI.

## MATERIALS AND METHODS

### Dataset generation

A planar, two-segment model of an upper limb with shoulder and elbow joints (Fig. 1a) was implemented in Simulink (MathWorks, MATLAB R2025a, Natick, MA). Each segment was represented as a uniform-density rigid cylinder; segment lengths and masses were scaled to a nominal adult (75 kg, 1.75 m) using standard anthropometric fractions (Winter, 2009). Segmental motion was described with coupled EOM, 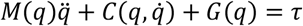, where joint angles, velocities, and accelerations 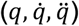 modulate posture-dependent inertia matrix (*M*), Coriolis and centrifugal forces (*C*), and gravity vector (*G*).

**Figure 1.**
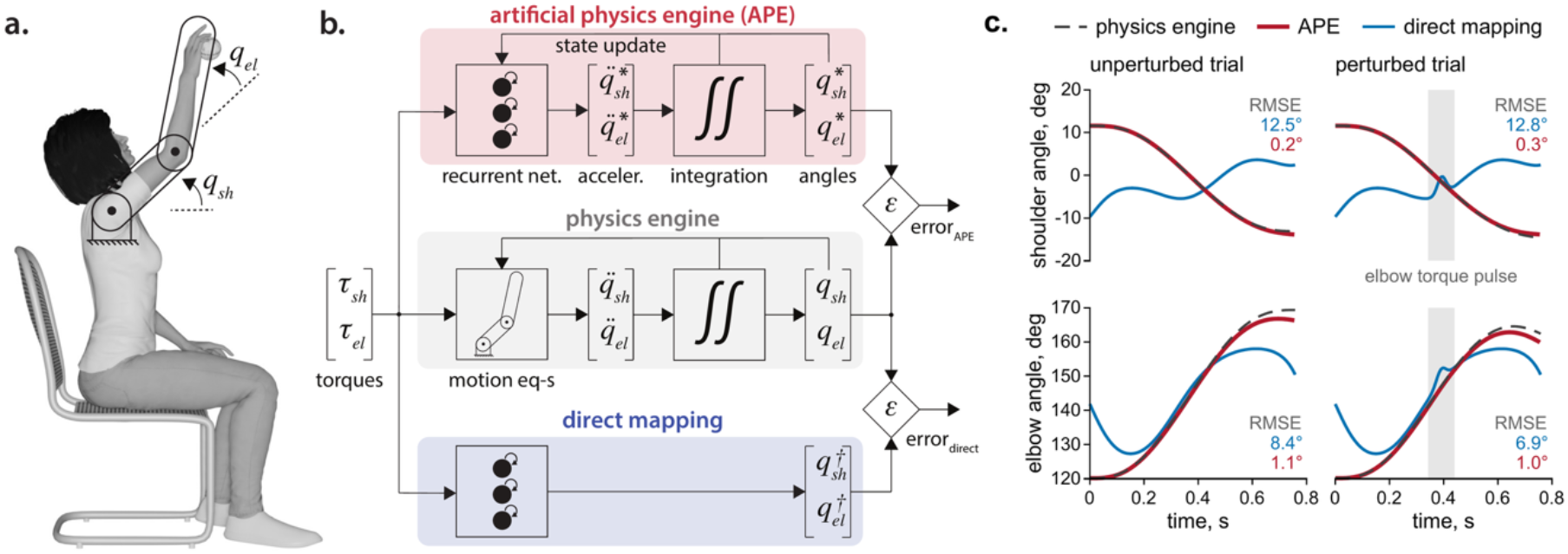
Comparison of the artificial physics engine and a direct mapping model in two-joint arm movements. (**a**) Schematic of the planar two-segment arm model used in simulations, representing shoulder (*q*_*sh*_) and elbow (*q*_*el*_) joint angles in the sagittal plane. Angles are referenced relative to an extended arm posture (defined as 0°, equivalent to 180° in conventional coordinates). (**b**) Block diagrams illustrating the three modeling approaches. The ground-truth physics engine (middle, gray) computes joint kinematics by integrating accelerations derived from the EOM. The APE (top, red) employs an RNN to learn the mapping from the current torques and previous kinematic state to instantaneous joint accelerations, which are then externally integrated to predict the movement trajectory. In contrast, the direct mapping model (bottom, blue) uses an RNN to map the torques directly to joint angles without an explicit integration step. (**c**) Representative single-trial predictions of shoulder and elbow angles during an unperturbed trial (left) and a trial with an external torque perturbation (right). The APE’s predictions (red lines) closely track the ground-truth trajectories from the physics engine (dashed gray lines), while the direct mapping model (blue lines) shows substantial error. The gray band marks the torque pulse applied at the elbow, and the values give the RMSE of each model over the movement shown (blue, direct mapping model; red, APE).

Motion of the arm in a sagittal plane, subject to gravity, was simulated between three sets of targets, with one target in common, arranged radially on a circle within the reachable workspace (radius = 0.5 × forearm length). The training, testing, and validation movement sets were organized as start– end pairs between all target vertices (*n*_*train*_ = 12 resulting in 132 movements; *n*_*test*_ = 10 resulting in 90 movements, and *n*_*valid*_ = 7 resulting in 42 movements). Each movement was a straight-line endpoint path with bell-shaped velocity (Morasso, 1981) specified in Cartesian space and converted to joint space using inverse kinematics. We solved the inverse dynamics problem to compute shoulder and elbow torques using analytically derived joint angles and the velocities and accelerations obtained from them by finite differences. Backward Euler with a simulation step size *Δt* of 1 µs provided sufficient temporal resolution to ensure numerical stability and accuracy. To generate movements with imposed brief perturbations, we added a Gaussian pulse (curtailed to a 3-sigma kernel, set to 50-100 ms) to either of the joint torques, randomizing the joint and direction. The changed trajectory was then computed by numerically integrating the EOM with a temporal resolution of 1 µs. The pulse amplitude was set to 5–15% of the peak-to-peak range of that joint’s unperturbed torque in the corresponding movement. The perturbation onset was drawn uniformly from 20–80% of the movement duration. The perturbed movements between the 7 validation targets were included in the training process *only* as validation trials, used to monitor training progress and detect potential overfitting, but *not* to update weights. The dataset with perturbed movements generated for the 10 test targets was used to evaluate whether the trained RNN models appropriately embedded the segmental dynamics accounting for the interaction torque in movements with perturbations. All signals were resampled by linear interpolation to 0.1 ms for training and testing.

### Model training

We used three models in this study, two RNNs and a standard mechanical model of a two-segment system, described above (Fig. 1b). The APE (top panel) was trained to predict instantaneous joint accelerations from the current torques and the preceding kinematic state, computed on the previous iteration. For each instance (*t*), the input consisted of current shoulder and elbow torques (*τ*_*s*_, *τ*_*e*_) paired with the prior joint angles and velocities 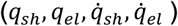, at time *t* − Δ*t*, yielding a 6-dimensional input vector. The target output was the predicted acceleration vector 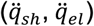 at *t*. The network was configured as a single long short-term memory (LSTM) layer (16 hidden units) that processes each time step independently and carries no internal state between steps, followed by a linear 2-unit regression head. Input normalization was disabled, and L2-norm gradient clipping was set to 1. The second RNN, a direct mapping model (bottom panel), was trained to map joint torques directly to the current joint angles, omitting kinematic inputs. This network had the same architecture as the first network. Both RNNs were trained with the Adam solver to minimize root-mean-square error (RMSE) over their outputs, joint accelerations for the APE and joint angles for the direct mapping model (initial learning rate 3×10^-3^, with a piecewise schedule reducing the rate by a factor of 2/3 every 50 epochs), using mini-batches of 8192 samples and shuffling the data within each epoch for up to 150 epochs. Performance on a validation set was monitored, and the model weights corresponding to the lowest validation RMSE were retained.

### Simulating dynamics with neural networks

After training, the networks were used in their respective models (APE or direct mapping) to predict joint angles across tests in unperturbed and perturbed movement conditions. Movements spanned all ordered start-end pairs among the 10 test targets, and none of these movements was seen during training. For the APE, the prediction for a given movement was initialized with its initial kinematic state (the first sample of the reference data). At each subsequent step, the embedded network used the current torques and the kinematic state from the previous iteration step to predict the instantaneous joint accelerations. Joint velocities and angles were then updated using explicit Euler integration (*Δt* = 0.1 ms) of the predicted accelerations. This ‘predict–integrate’ cycle, in which the newly computed state becomes the input to the next step, was repeated until the end. Except for the first state-initialization step, reference kinematic data were *not* used in this process. State estimation relied solely on numerically integrating the network-predicted accelerations, driven by the torque stream. For the direct mapping model, the instantaneous torques were the only inputs used to predict joint kinematics. Predictions were generated independently for each point over the entire movement without explicit numerical integration.

The APE solved an initial value problem in which the kinematic state (joint angles and velocities) was the only quantity carried between time steps, because the network processed each time step independently. This carry-over of the computed state (state update, Fig. 1b) reproduced the computation chain of standard numerical integration, and the ground-truth physics engine, which generated the perturbed trajectories, performed the corresponding update with the EOM in place of the network. The state update was the only loop in the APE, closed by the explicit Euler integrator of the decoupled architecture, and no feedback from the reference kinematics corrected the state. Every acceleration error therefore propagated into all subsequent velocities and angles, and the RMSE between simulated and reference joint angles quantified the error accumulated over the entire movement.

### Statistical analysis

The normality of the RMSE distributions for each model and simulated movement type (perturbed/unperturbed) was assessed using the Lilliefors test. Because all data distributions were nonnormal, a nonparametric Kruskal-Wallis test was used to assess whether significant differences existed among the groups. Pairwise comparisons were conducted using a post-hoc test with Dunn-Sidak correction to identify which specific groups differed from one another. All statistical analyses were performed in the Statistics and Machine Learning Toolbox (MathWorks), with the significance level set at α = 0.05.

## RESULTS

In this study, we show that decoupling state-dependent dynamics from numerical integration enables neural networks to simulate forward dynamics. Monolithic, direct-mapping models conflate these computational steps, increasing representational complexity and failing to approximate perturbed movements. By contrast, the decoupled architecture of our APE model isolates the equations of motion and delegates state integration to an external solver. This modular approach reduces prediction error compared with the monolithic baseline. The APE generalizes to novel initial conditions and unseen perturbations while exhibiting emergent intersegmental dynamics, suggesting that the model internalizes the system’s physics rather than memorizing trajectories.

### Direct mapping

Under matched training protocols, the direct model, with end-to-end mapping from joint torques to joint angles, yielded substantially higher prediction errors than the modular APE model. As illustrated by examples of both unperturbed and perturbed movements (Fig. 1c), the direct model performed poorly at generalizing across trials and failed to associate input torques with the corresponding movement evolution. Lacking initial conditions, it could not associate initial torque values with the appropriate starting state. This limitation arises in part because the same instantaneous torques can correspond to different joint angles, depending on the kinematic state.

Across 90 unperturbed trajectories, the RMSE (mean ± s.d.) was 15.74 ± 6.17° for the shoulder and 12.04 ± 3.96° for the elbow (Fig. 2a, blue). Under external perturbations, the network performed similarly poorly, yielding RMSE values of 15.71 ± 6.17° for the shoulder and 12.20 ± 4.07° for the elbow (Fig. 2b, blue). These errors did not differ significantly from those in the unperturbed condition (p > 0.05, post hoc pairwise comparisons with the Dunn–Sidak correction following the Kruskal– Wallis test).

**Figure 2.**
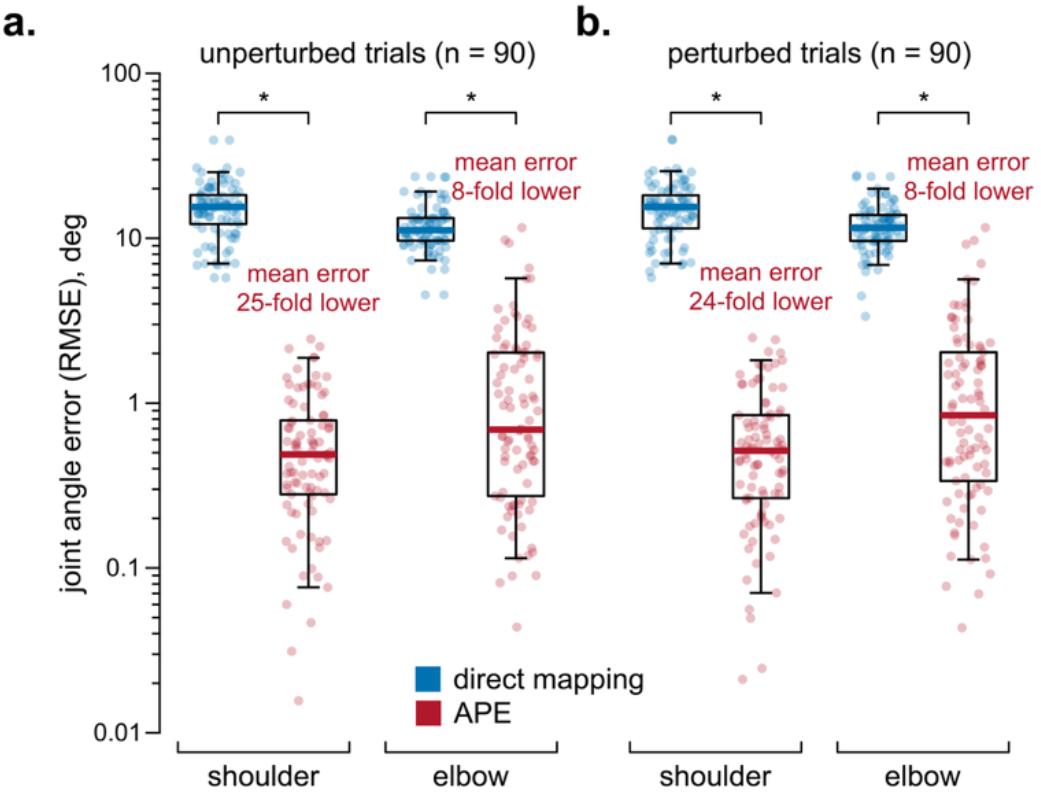
The direct mapping model underperforms relative to APE, the modular model, as illustrated by predicted shoulder and elbow angle RMSE across all test trials. Each point represents a single trial. Boxes show the median and quartiles, and whiskers the 5th and 95th percentiles, on a logarithmic axis. (**a**) The comparison between the direct model (blue) and APE (red) is shown for 90 unperturbed trials. The APE reduced the mean prediction error by approximately 25-fold for the shoulder and 8-fold for the elbow compared with the direct mapping model. (**b**) This high level of accuracy was maintained across 90 perturbed trials, with APE performance not significantly different from that in the unperturbed condition. In contrast, the direct mapping model exhibited consistently high errors that also did not differ significantly between unperturbed and perturbed conditions. Asterisks (*) indicate significant differences between the two models (p < 0.01, post hoc pairwise comparisons with the Dunn–Sidak correction following the Kruskal–Wallis test).

### Decoupling physics from integration

The APE model was configured to treat the mapping as an initial value problem and yielded significantly more accurate and robust predictions than the direct mapping approach. Figure 1c shows representative examples where APE predictions (red) closely tracked reference trajectories during both unperturbed and perturbed trials. The APE achieved mean RMSE values of 0.64 ± 0.53° for the shoulder and 1.56 ± 2.10° for the elbow in unperturbed trials, representing a 25-fold and 8-fold reduction in error compared to the direct mapping approach (Fig. 2a; p < 0.01, post-hoc pairwise comparison with Dunn-Sidak correction). The model maintained high accuracy when external perturbations were applied, with error values of 0.66 ± 0.56° for the shoulder and 1.62 ± 2.11° for the elbow that were not significantly different from unperturbed conditions (Fig. 2b; p > 0.05, post-hoc pairwise comparison with Dunn-Sidak correction). This stability, maintained even when faced with new initial conditions and external perturbations (perturbed movements were never used to update the weights), suggests that the model had internalized the system’s EOM.

### Predicting complex intersegmental dynamics

To test whether APE internalized the EOM, we evaluated its prediction of intersegmental dynamics during untrained, perturbed trajectories. Figure 3 illustrates a representative trial. A transient elbow perturbation (bottom panel, gray shaded region) induces a compensatory shoulder acceleration via inertial coupling. In learning the forward mapping 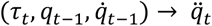, this interaction is governed by the mass matrix *M*(*q*) ∈ ℝ^*n*×*n*^. For the two-link system, the mass matrix is defined as

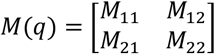

**Figure 3.**
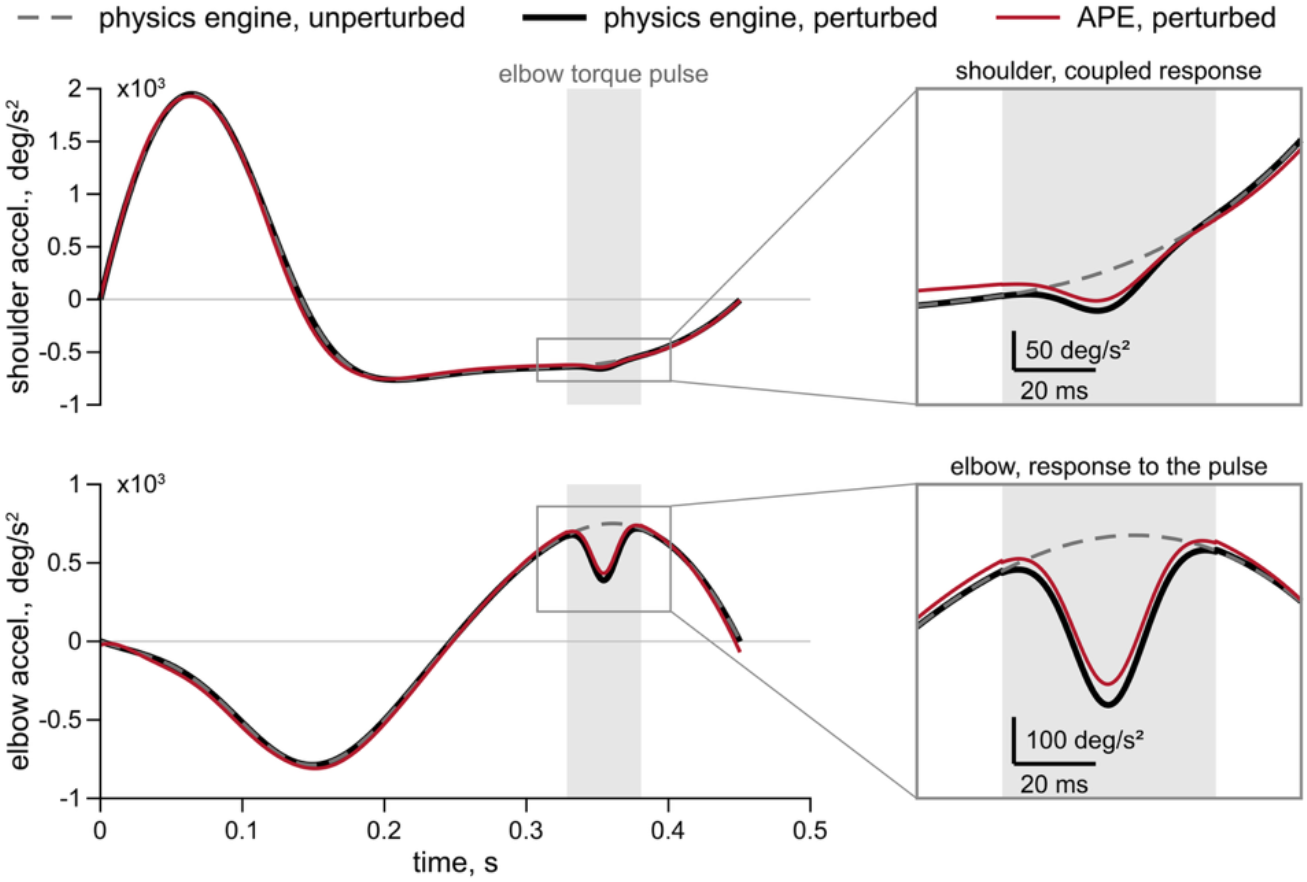
The APE predicts intersegmental dynamics. Shown is a representative trial demonstrating the APE’s ability to predict the mechanical coupling between joints. The plots show shoulder (top) and elbow (bottom) accelerations for the physics engine without the perturbation (dashed gray line), the physics engine with the perturbation (solid black line), and the APE with the perturbation (solid red line). A transient torque perturbation was applied to the elbow (bottom, gray shaded region), causing a coupled, compensatory acceleration at the shoulder due to interaction torques (top, gray shaded region). The APE’s prediction captures about 80% of this compensation (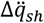, the difference between the solid black and dashed gray lines in the upper expanded view). Gray frames show the interval around the perturbation on an expanded scale, with scale bars for time and acceleration.

Consequently, the shoulder torque 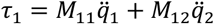, depends directly on elbow acceleration (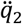). The APE reproduced about 80% of this coupled shoulder response (−40 against −49 deg/s^2^; top panel, expanded view) despite training exclusively on unperturbed movements, suggesting that the network embedded the physical equations governing the system.

## DISCUSSION

In this study, we demonstrate that decoupling the process of integration from the equations of motion, rather than simulating a direct trajectory mapping, provides a more computationally tractable formulation for forward dynamics learning. We hypothesized that the predictive instability of monolithic neural networks in forward dynamic simulations stems from conflating two distinct computational steps: (1) approximating physical laws and (2) performing numerical integration. To resolve this, we developed an artificial physics engine (APE) that explicitly decouples these processes. Using a shallow RNN to approximate the instantaneous accelerations of a limb model, and delegating numerical integration to a dedicated external solver, the APE reduced prediction error 25-fold at the shoulder and 8-fold at the elbow relative to traditional monolithic networks. This physics-embedded architecture not only generalized to unencountered initial conditions and perturbations but also reproduced complex, interaction-torque-driven intersegmental dynamics, consistent with an embedding of the underlying mechanical laws. *Ultimately, these findings support the broader principle that the computational tractability of learning movement dynamics depends not only on network capacity, but also on the representation in which the problem is posed*.

With network architecture and training protocol matched, posing forward dynamics as state evolution in place of direct mapping reduced mean shoulder RMSE from 15.74° to 0.64° and mean elbow RMSE from 12.04° to 1.56° (Fig. 2a). Control theory distinguishes these two formulations (Kalman, 1963), and the direct mapping model implements the input-output mapping without explicit internal state that is common in movement decoding (Glaser et al., 2020; Mathis et al., 2024), whereas the APE implements the state-evolution equation. The EOM determine accelerations uniquely from the torques and the kinematic state, and the network within the APE receives both as inputs. The direct mapping model must instead absorb the initial state and the integration over the torque history into a function of the instantaneous torques, and its trajectories started away from the reference posture (Fig. 1c). The APE, with one 16-unit LSTM layer, is a representative implementation that suffices for stable, accurate simulation of a two-segment arm under unseen perturbations, and optimization of architecture and training may refine this accuracy further. This proof of principle shows that the decoupled architecture gives HMI design a forward model that simulates an entire movement from the torque stream and a single initial state.

The primary limitation of our study is the reliance on a simple two-segment mechanical model to represent the upper arm and forearm movement within a vertical plane. While this model does not provide sufficient detail to simulate fine motor tasks, such as hand-object manipulation, finger coordination, and detailed contact mechanics, it is sufficient to probe whether a learning system captures coupled multi-joint dynamics. Two-segment models remain valuable because they incorporate multi-segment coupling; their dynamics include inertial, Coriolis, and centrifugal interaction terms absent in single-joint movements. This level of modeling is often sufficient to determine whether a method captures the essential properties of a coupled articulated structure and is commonly used to represent human biomechanics during planar reaching, describing how torque terms, including intersegmental interaction terms, shape movement (Gritsenko et al., 2009; Hollerbach and Flash, 1982). Similar mechanical complexity is also relevant to gross locomotor modeling that captures coordination and energetics (Darici and Kuo, 2022; Faraji and Ijspeert, 2017; Lin et al., 2020; Saunders et al., 1953; Taga et al., 1991), with applications in prosthetics. The 2-DOF model employed in this study represents the simplest mechanism suitable for testing embedded multijoint physics.

### Biomimetic learning of system dynamics via prediction error

The APE’s acquisition of the system dynamics of the double pendulum mimicked the hypothesized acquisition of bodily dynamics by the internal forward model within the central nervous system. Such a process is driven by sensory prediction errors, computed by comparing expected and actual outcomes. In the biological motor system, the forward internal model processes the efference copy of the motor command and generates a prediction of the expected sensory consequences, such as the dynamics of the effector (Wolpert and Flanagan, 2001); our artificial engine used an analogous process, in which the RNN processed the torque command and generated a prediction of the pendulum’s dynamics, with the actual dynamics reported by the physics engine serving as the ‘sensory’ feedback. The discrepancy between predicted and sensory feedback constituted the sensory prediction errors. These errors are widely believed to drive synaptic modification within the biological circuits forming the forward internal model (Shadmehr et al., 2010). Analogously, in the neural network within the APE, prediction errors were used to tune the weight matrices via a learning rule. This error-driven tuning enabled the APE to accurately respond to unseen external perturbations, functionally replicating the entrained adaptive responses well-documented in biological motor control studies, for example, force-field learning paradigms (Shadmehr and Mussa-Ivaldi, 1994; Thoroughman and Shadmehr, 2000).

### Overcoming learning and simulation challenges

A primary obstacle to designing accurate and stable noninvasive HMIs is the non-stationary relationship between EMG signals and movement (see Scheme and Englehart, 2011 for review), as these signals are continuously modulated by intrinsic and extrinsic factors in real-time. For conventional data-driven models that learn this direct mapping, non-stationarity poses a critical problem known in deep learning as *covariate shift*, where the statistical distribution of the input data in real-world use differs from the distribution of the data used to train the model (Vidovic et al., 2016, 2014). This shift results in low, hard-to-learn performance that may degrade as the interface parameters change, leading to the need for frequent recalibrations. The evidence of out-of-sample prediction of interaction torques with APE indicates that the network simulates a causal, physical relationship between applied forces and resulting motion.

However, the current APE implementation, which was focused on high physical precision, has not achieved real-time performance (using a standard personal computer) with the implementation limited to the choice of a fixed-step explicit Euler integrator to propagate the system’s state. To ensure numerical stability and accuracy when simulating the complex, nonlinear, and sometimes stiff dynamics of the two-segment arm, a small, fixed time step (*Δt* = 0.1 ms) was necessary, but incompatible with the strict time budgets of interactive applications like prosthetic control or virtual reality. The typical goal for a full simulation and rendering cycle should be less than 33 ms to maintain the illusion of smooth motion (Dinev et al., 2018), with the physical simulation loop latency under 16 ms (Erez et al., 2015), and even shorter latencies for the accurate computations of muscle models. An important future direction for this work is, therefore, to transition the APE from a fixed-step to an adaptive step-size integration scheme. Such predictive allocation of computational resources is computationally biomimetic, since biological motor control is inherently dynamic, allocating neural resources to meet task demands (Todorov, 2004).

An alternative to the numerical solver is an architecturally suitable neural network, such as a residual neural network. Characterized by skip connections of the form *h*^*l*+1^ = *h*^*l*^ + *f*(*h*^*l*^), where *h*^*l*^ is the hidden state at layer *l* (representing a discrete step of numerical integration) and *f* is a learned transformation modeling system dynamics, a residual network is mathematically equivalent to the forward Euler discretization of an underlying ordinary differential equation (also see Chen et al., 2019; He et al., 2015). Hence, the APE could, in principle, be formulated as a two-stage neural architecture: (1) a recurrent network to approximate the EOM, and (2) a residual network trained to perform the temporal integration. Whether a *fully* neural, two-stage APE offers practical advantages over the current hybrid approach for HMI applications, which combines a neural network approximation of the motion equations with a classical integrator, remains an open question.

In this study, we propose a novel framework for embedding forward dynamics within shallow neural networks that encode the formulation of mechanics separately from its integration. We tested our hypothesis that the decoupled architecture can learn the multisegmental physics of interactions by generalizing observed movement, without the explicit exposure to perturbations. The accurate and stable representation of dynamics in this framework establishes a viable technical path for the broader strategy of embedding internal forward models in advanced HMIs.

## DATA AND CODE AVAILABILITY

The data underlying the figures are available at https://github.com/SerhiiBahdas/ms-frw-ape. Simulated training and validation datasets can be obtained from the corresponding author upon reasonable request. Scripts for regenerating figures, along with run instructions, are provided in the same repository. Code for simulating arm biomechanics, training the networks, and running the APE is available from the corresponding author upon reasonable request.

## ACKNOWLEDGMENTS

This work was supported by the Air Force Office of Scientific Research (Award No. FA9550-24-1-0208), the West Virginia Foundation Distinguished Doctoral Scholarship Program, the West Virginia University Center for Foundational Neuroscience Research and Education, and the West Virginia University Stroke and Alzheimer’s Disease-Related Dementias (ADRD) T32 Program. S.B. also acknowledges the NeuroMechatronics Lab for their valuable advice and comments.

